# Climatic and biogeographic processes underlying the diversification of the pantropical and early divergent angiosperm family Annonaceae

**DOI:** 10.1101/2023.08.08.549183

**Authors:** Weixi Li, Runxi Wang, Ming-Fai Liu, Ryan A. Folk, Bine Xue, Richard M.K. Saunders

**Affiliations:** Division of Ecology & Biodiversity, School of Biological Sciences, The University of Hong Kong, Hong Kong, China; Department of Biological Sciences, Mississippi State University, Starkville, Mississippi, USA; Zhongkai University of Agriculture and Engineering, Guangzhou, China

**Keywords:** diversification rate shift, global temperature, biogeographical events, climatic niche evolution, diversification process

## Abstract

**Aim:** Tropical rainforests harbour the richest biodiversity among terrestrial ecosystems, but few studies have addressed underlying processes of species diversification in these ecosystems. We use the pantropical and early divergent flowering plant family Annonaceae as a model system to investigate how abiotic factors such as climate and biogeographic events contribute to the diversification process and lead to its high diversity across a long evolutionary history.

**Location:** Tropics and subtropics

**Taxon:** Annonaceae

**Methods:** A super-matrix was constructed for 835 taxa (34% of Annonaceae species), based on eight chloroplast regions. To understand the patterns of diversification, we reconstructed climatic niche evolution and historical biogeographical events, and tested their association with diversification rates.

**Results:** The analysis of temperature-dependent models in Annonaceae lineages provides strong support for the significant influence of global temperature on net diversification and accumulation of species diversity. The pattern of lineage accumulation in the initial radiation is better aligned with the “museum model,” followed by later accumulation consistent with the “recent cradle model” from the late Oligocene to the present. The increase in the diversification rate of the family (around 25 Ma) lags behind the accumulation of niche divergences (around 15 Ma). Biogeographic events are related to only two of the five diversification rate shifts detected. While no direct relationship to shifts in the diversification rate was uncovered, shifts in niche evolution appear to be associated with increasingly seasonal environments.

**Main Conclusions:** Global temperature plays a crucial role in driving recent rapid diversification in the Annonaceae. Our study challenges the prevailing assumption of the “museum model” alone and proposes instead a transition from the “museum model” to the “recent cradle model” during the diversification history of the family. However, our findings do not support the direct correlation of any particular climatic niche shifts or historical biogeographical events with shifts in diversification rate. Instead, Annonaceae diversification can lead to later niche divergence as a result of increasing interspecific competition arising from species accumulation. The evolutionary direction of niche shifts furthermore provides insight into the future expansion of Annonaceae into temperate regions. Our results highlight the complexity of the diversification process in taxa with long evolutionary histories, indicating that identifying isolated driving factors is simplistic and inadequate for explaining the observed patterns. Further comprehensive analyses of range evolution are necessary to delve deeper into the interplay among key opportunities, key innovation, and species diversification.

## 1. Introduction

Tropical rainforests occupy 7% of land surface and greatly contribute to the species diversity of forest ecosystems (Wilson, 1988; Whitmore, 1998; Hill & Hill, 2001; Lomolino et al., 2010). Tropical rainforest biomes face the threat of extinction during the ever-changing climate (Temperton et al., 2019). The processes and mechanisms underlying the diversification of species in tropical rainforests remain unclear, hampering the assessment of how tropical rainforest ecosystems respond to global change and hence impeding the development of conservation priorities in the face of future climate change.

Immigration, speciation, and extinction events drive diversification. Current evidence suggests that immigration into tropical rainforests is very limited (Donoghue & Edwards, 2014), however, implying that speciation and extinction are perhaps the key to species diversification of rainforests. The path from speciation and extinction to modern-day diversity depends on age and rates: in other words, the relationship between the very high species richness of rainforests and evolutionary rates in these biomes is not straightforward. Several models have been developed to predict lineage accumulation patterns: in the “ancient cradle model”, lineages accumulate with a high initial speciation rate and then decelerate towards the present, with a low overall extinction rate; in the “recent cradle model”, in contrast, lineage accumulation is rapid towards the present; finally in the “museum model”, lineages accumulate at a constant rate, with low extinction overall and with diversity primarily reflecting time since origin (Wallace, 1878; Gentry, 1982; Richardson et al., 2001; Zachos et al., 2001; Davis et al., 2005; Morley et al., 2007; Stebbins, 2013). The cradle and museum hypotheses are not mutually exclusive: since speciation-extinction dynamics change through time, tropical rainforest taxa can adopt a museum regime for one lineage at a specific time and a cradle regime at another time.

Multiple factors have been proposed as drivers of diversification. On a geological timescale, global climate changes have repeatedly been shown to significantly influence diversification (Erwin, 2009), with recent botanical examples including *Primulina* (Gesneriaceae; Kong et al., 2017), Saxifragales (Folk et al., 2019), rosids (Sun et al., 2020), Asteraceae (Zhang et al., 2021), *Scleria* (Cyperaceae; Larridon et al., 2021), and *Oreocharis* (Gesneriaceae; Kong et al., 2022). Global temperature has fluctuated dramatically throughout Earth’s history (Hansen et al., 2006; Zachos et al., 2001; Rahmstorf et al., 2017) and is considered to play a crucial role in shaping the fate of clades and the biomes that they are adapted to. Two contrasting evolutionary mechanisms might be invoked to explain how taxa respond to global temperature variation: “key innovations” involving intrinsic biotic factors, such as morphological or behavioral changes, can enable lineages to overcome abiotic stressors and enable novel niche invasions (Larridon et al., 2021; Moore and Donoghue, 2007). By contrast, “key opportunities” involve pre-existing adaptations that enable lineage survival in a changing abiotic environment, including climatic variation through time and biogeographical events (Moore and Donoghue, 2009; Donoghue and Sanderson, 2015). The essential difference between these is that a key innovation narrative implies agency on the part of species adapting to changing situations, whereas key opportunity narratives invoke species that are “in the right place at the right time” with traits preadapted for changing conditions. Some studies have highlighted the importance of climate niche evolution in the diversification of animals (Reis et al., 2018; Kozak & Wiens, 2010; Li & Wiens, 2022). While Liu et al. (2020) have demonstrated similarities in the evolution of climatic niches between animals and plants, because of the restricted migration abilities of plants there are relatively few studies that explore the relationship between macroevolution and biogeography in tropical plant species. Rather than being hypotheses in conflict, numerous studies suggest key innovations and key opportunities can synergistically play a role in species diversification (Moore & Donoghue, 2007; Larridon et al., 2021). Integrating temperature- and state-dependent diversification models and conducting comparative analyses would help disentangle the association between abiotic factors and tropical diversity and distinguish among hypotheses explaining the distribution of this diversity.

The family Annonaceae (Magnoliales) represents an ideal system for understanding species diversification in tropical rainforest ecosystems. The family originated in the middle Mesozoic (112.6–89 Ma; Xue et al., 2020), and is a large and highly diverse pantropical lineage, comprising approximately 2440 species, widely distributed in lowland forests (Couvreur et al., 2012). Previous studies have suggested that lineages within Annonaceae accumulated through stable diversification rates over time (Couvreur et al., 2011), implying a gradual increase in species consistent with the museum model of diversification (Couvreur et al., 2011; Stebbins, 1974). The limited taxon sampling in the previous study (4.8% of species: Couvreur et al., 2011), however, is likely to have resulted in an underestimation of diversification rate. More recent research with improved sampling (34.2% of species) has challenged the museum model interpretation, suggesting instead an accelerated diversification in the Annonaceae from ca. 25 Ma, both at the family and major clade levels (Xue et al., 2020). While certain biotic factors in the Annonaceae, such as pollinator trapping, androdioecy, and single-seeded monocarp fragments, are possibly correlated with high diversification rates (Xue et al., 2020), the contribution of abiotic factors, including biogeography and climate changes, to the rapid diversification rate increase has yet to be investigated. The combination of incomplete sampling in previous work on the family (Couvreur et al., 2011), uncertainty about the timing and drivers of diversification, and the paucity of studies concerning the evolution of pantropical rainforests suggests that our overall framework for understanding tropical diversification is incomplete. While a recent study suggests that the recent cradle model is applicable (Xue et al., 2020), a more comprehensive sampling and phylogenetic construction are needed for revealing the diversification dynamics of Annonaceae and strengthening their status as a model for tropical rainforest diversification.

Correlations between key opportunities (biogeographical events) or key innovations (niche or trait evolution) and diversification rate shifts in other plant families are not always deterministic (Moore & Donoghue, 2000; Larridon et al., 2021; Stull et al., 2021), which highlights the need to consider multiple factors simultaneously. While a recent study by Wang et al. (2021) revealed a correlation between the rate of climate niche evolution and diversification in Annonaceae, the specific sequence and interplay between key innovation and key opportunities in relation to diversification remain obscure and are needed to more fully investigate tropical plant diversification models. Our research addresses this knowledge gap, and explores the following hypotheses: (1) Is species accumulation in the Annonaceae appropriately characterized by the recent acceleration model? (2) Does temperature drive diversification of the family? (3) Do niche and/or range evolution play a role in diversification of the Annonaceae—and if so, what are the sequence of those events, and do they act independently or synergistically?

## 2. Methods

### 2.1 DNA data sampling

The alignment file, comprising 916 Annonaceae ingroup and seven outgroup taxa following concatenation of eight chloroplast genes or gene spacer regions, was obtained from Xue et al. (2020). This data matrix is the best sampled to date. A pruned alignment file was generated with 835 tips (after excluding outgroup taxa); undescribed and unidentified species were also removed to eliminate potential misidentifications or uncertainties.

### 2.2 Divergence time estimation

In this study, Maximum Likelihood (ML) analyses based on the 835-taxon tree with 1000 rapid bootstrapping were conducted using RAxML v.8.2.12 (Stamatakis, 2014), performed on the CIPRES Science Gateway v.3.3 online server (https://www.phylo.org/portal2/home.action). The analyses utilized the GTRGAMMA evolution model, with the simultaneous searching algorithm employed to generate the best ML tree. Bootstrap resampling was applied with 1000 replicates to assess topological support. Divergence times were estimated using the penalized likelihood method implemented in the treePL software (Smith & O’Meara, 2012) due to its efficiency in handling large datasets compared to other tools such as BEAST. The input trees consisted of the 1000 ML bootstrap trees obtained from RAxML, with branch lengths preserved. The root age of the phylogenetic tree was fixed at 137 Ma based on the divergence time between Magnoliales and Laurales as recovered by Foster et al. (2017). This age is also supported by the presence of angiosperm crown group pollen grain fossils from the Hauterivian (Early Cretaceous, 136.4–130 Ma: Hughes, 1994, Friis et al., 2010). For other calibrations, the Annonaceae crown node was set between 112.6 and 89 Ma, while the Magnoliineae crown node was placed between 125 and 112.6 Ma, following the CS3 calibration scheme (Thomas et al., 2015; Thomas et al., 2017).

### 2.3 Patterns of lineage accumulation

Semi-logarithmic lineages-through-time (LTT) plots were generated based on the 835-taxon tree to visualize the pattern of species richness accumulation. The LTT plots were created using the laser package v.2.2 (Rabosky, 2006) in R. LTT plots were generated separately for two levels: (1) the family-level LTT plot, representing the overall species richness pattern for the entire family; and (2) LTT plots for the four subfamilies within Annonaceae, allowing a more detailed examination of species richness dynamics within specific subgroups. To obtain robust estimates, 1000 phylogenetic trees were used in the analysis. For each tree, a mean LTT plot was generated, representing the average species richness through time. Additionally, a 95% confidence interval (CI) was calculated to assess the uncertainty associated with the estimates.

### 2.4 Paleotemperature-dependent diversification

In this study, a global temperature-dependent diversification analysis based on 835-taxon tree was conducted using the paleotemperature Cenozoic curve published by Zachos et al. (2008). The fit_env function in the R-package RPANDA v.1.1 (Morlon et al., 2016) was employed to evaluate the best diversification model. Eight models were fitted to environmental data, as follows: (1) BEnvVar EXPO: speciation rates vary exponentially with environmental data, while the extinction rate is fixed at zero (unparameterized); (2) BEnvVar DCST EXPO: speciation rates vary exponentially with environmental data, while the extinction rate remains constant; (3) BCST DEnvVar EXPO: extinction rates vary exponentially with environmental data, while the speciation rate is constant; (4) BEnvVar DEnvVar EXPO: both speciation and extinction rates vary exponentially with environmental data; (5) BEnvVar LIN: speciation rates vary linearly with environmental data, while the extinction rate is fixed at zero; (6) BEnvVar DCST LIN: speciation rates vary linearly with environmental data, while the extinction rate is constant; (7) BCST DEnvVar LIN: extinction rates vary linearly with environmental data, while the speciation rate is constant; and (8) BEnvVar DEnvVar LIN: both speciation and extinction rates vary linearly with environmental data. Eight time-dependent birth-death models and two constant rate models were furthermore included in the model set for comparison. The constant rate models were: (9) BCST: pure birth model with constant speciation rate but extinction rate fixed at zero; and (10) BCST DCST: both speciation and extinction rates vary by a constant rate. Comparison of these models enables the identification of the best-fitting model that describes the relationship between temperature and diversification rates. The results were plotted using RevBayes (Höhna et al., 2016) based on the best fitting model estimated by RPANDA.

### 2.5 Biogeography

The distribution data for the study were obtained from the Global Biodiversity Information Facility (GBIF; http://www.gbif.org), with additional locality data sourced from iDigBio (https://www.idigbio.org/portal/search). The dataset was manually checked and cleaned using “CoordinateCleaner” (Zizka et al., 2019) in R, with a total of 728 species with specific locations. Therefore, species without specific location data were pruned from the 835-taxon tree resulting a tree of 728 species for biogeographical analysis.To infer the ancestral areas of extant species and reconstruct dispersal history, all extant species were assigned to six geographical regions based on palaeogeographical reconstructions (Couvreur et al., 2011) and current Annonaceae distribution data: (A) Southeast Asia, west of Huxley’s Line; (B) Southeast Asia east of Huxley’s Line, northern Australia, and Pacific islands; (C) Continental Africa; (D) Madagascar; (E) North/Central America; and (F) South America. The maximum size of reconstructed ancestral areas was set to six regions. Ancestral area reconstruction was performed using BioGeoBears (Matzke, 2013) under four models: Dispersal-extinction-cladogenesis (DEC), and Dispersal Vicariance Analysis (DIVALIKE), and their +j variant (the jump dispersal parameter, allowing for “founder-event speciation”). The best-fitting model was determined based on the lowest corrected Akaike’s information criterion (AICc) values, enabling identification of the scenario that provides the best trade-off between model fit and complexity.

Biogeographical stochastic mapping (BSM) was conducted to estimate the number and type of geodispersal events based on the best-fitting model. The frequencies and directions of these events were determined by calculating the mean and standard deviation (SD) from 50 BSMs. A time-stratification analysis was furthermore performed to estimate the number of dispersal events per 1 million years.

### 2.6 Climatic niche evolution

Bioclimatic data representing 19 temperature and precipitation measures were obtained from Worldclim (https://www.worldclim.org/) to investigate the evolution of bioclimatic variables and their potential association with speciation acceleration in Annonaceae. Mean values for each variable were calculated for each species, by using a dataset of 728 species, the same dataset as biogeography analysis, with valid bioclimatic data. Principal Component Analysis (PCA) was employed by using the function corrplot (Wei et al., 2017) in R to characterize the variation in climatic variables among species, thereby summarizing variance in the 19 variables into two-dimensional axes for downstream analysis. Mean PC scores were calculated for each species on each axis, providing a representation of their climatic niche. The R package “bayou” (Uyeda & Harmon, 2014) was used to infer climatic niche diversification rate shifts; this package utilizes a reversible-jump Bayesian method to fit the Ornstein-Uhlenbeck (OU) model. rjMCMC (reversible-jump Markov Chain Monte Carlo) sampling was run for 10^6^ cycles to achieve convergence, discarding the first 30% of samples as burn-in. Only one shift per branch was allowed, and shifts with posterior probabilities (PP value) above 0.5 were plotted to identify significant shifts in niche lability.

### 2.7 Time-dependent diversification

Diversification rates through time across the Annonaceae phylogeny based on the 728-taxon tree were estimated using the BAMM v.2.5 (Rabosky et al., 2014). BAMM uses a reversible-jump Markov chain Monte Carlo (rjMCMC) algorithm to explore the parameter space and estimate diversification rates under optimized “configuration” (sets of reconstructed shifts between diversification regimes with parameterizations within each). The MCMC analysis was run for 20,000,000 generations, with sampling every 1,000 generations. After discarding 10% of the output as burn-in, convergence was assessed by reference to effective sample size (ESS) values that exceed 200 for the likelihood and the inferred numbers of shifts using the coda package v.0.19-1 (Plummer et al., 2006) in R, to ensure adequate sampling from the posterior distribution. After the MCMC analysis, BAMMtools v.2.5 (Rabosky et al., 2014) was utilized to calculate the posterior distribution and to perform downstream analyses.

### 2.8 SSE analysis

The GeoSSE (Geographic State Speciation and Extinction) model (Goldberg et al., 2011) was employed to investigate the influence of geographical factors on the diversification of Annonaceae based on the dataset of 728 species. This model, implemented in the “diversitree” R package (Fitzjohn, 2012), allows for the examination of geographic range effects on diversification. SSE-type models have been shown to be vulnerable to Type-I error under certain conditions; to assess the potential impact of unmeasured factors on speciation and extinction in Annonaceae, HiSSE (Hidden State Speciation and Extinction) models (Beaulieu & O’Meara, 2016) were also utilized based on the dataset of 728 species. The best-fitting models were selected by comparing different models using likelihood-ratio tests under a chi-square distribution and Akaike information criterion (AIC). Four different scenarios were considered: (1) variation in dispersal parameters without range-dependent diversification; (2) the canonical GeoSSE model with range-dependent diversification; (3) the GeoHiSSE model with one hidden trait but without range-dependent diversification; and (4) the GeoHiSSE model with one hidden trait and range-dependent diversification. The model set therefore jointly tests for a diversification effect and for hidden states (diversification unexplained by measured states).

Quantitative analyses were conducted using the QuaSSE (Quantitative State Speciation and Extinction) model, implemented in the diversitree R package (FitzJohn, 2012), to examine the interplay between niche lability and diversification rates based on the dataset of 728 species. Two principal component (PC) axes values derived from 19 climatic variables were selected to estimate the evolution of speciation and extinction rates. The QuaSSE model incorporates a birth-death process, in which speciation (λ) and extinction rates (μ) are determined by a quantitative trait. We fitted three sets of QuaSSE models to explore different relationships between niche lability and speciation: (1) independent speciation rate; (2) linear relationship; and (3) sigmoidal relationship. The models were fitted both with and without the directional component (φ) to capture potential directional evolution of the trait (Fitzjohn, 2010). AIC values were used to determine the best-fitting models.

## 3. RESULTS

### 3.1 Pattern of lineage accumulation

The LTT plot of Annonaceae based on the 835-taxon tree, represented by the blue line in Fig. 1, shows a relatively constant species accumulation rate since its origin, with a recent radiation event at the late Oligocene. However, during the Cretaceous–Palaeogene (K/Pg) boundary period, the LTT plot of the entire family reveals a brief stasis period. Examining major sublineages, similar patterns are observed in the LTT plot of subfamily Annonoideae (Fig. S1: B), with a short stasis period at approximately 65 Ma and constant lineage accumulation until 30 Ma, after which diversification rate accelerated. Likewise, the LTT plot of subfamily Malmeoideae (Fig. S1: D) shows a fairly linear trend for a long period, followed by a rapid increase in diversification since approximately 25 Ma. Overall, species accumulation exhibits a prolonged period of stasis, followed by an acceleration of diversification starting from the late Oligocene.

**Figure 1.**
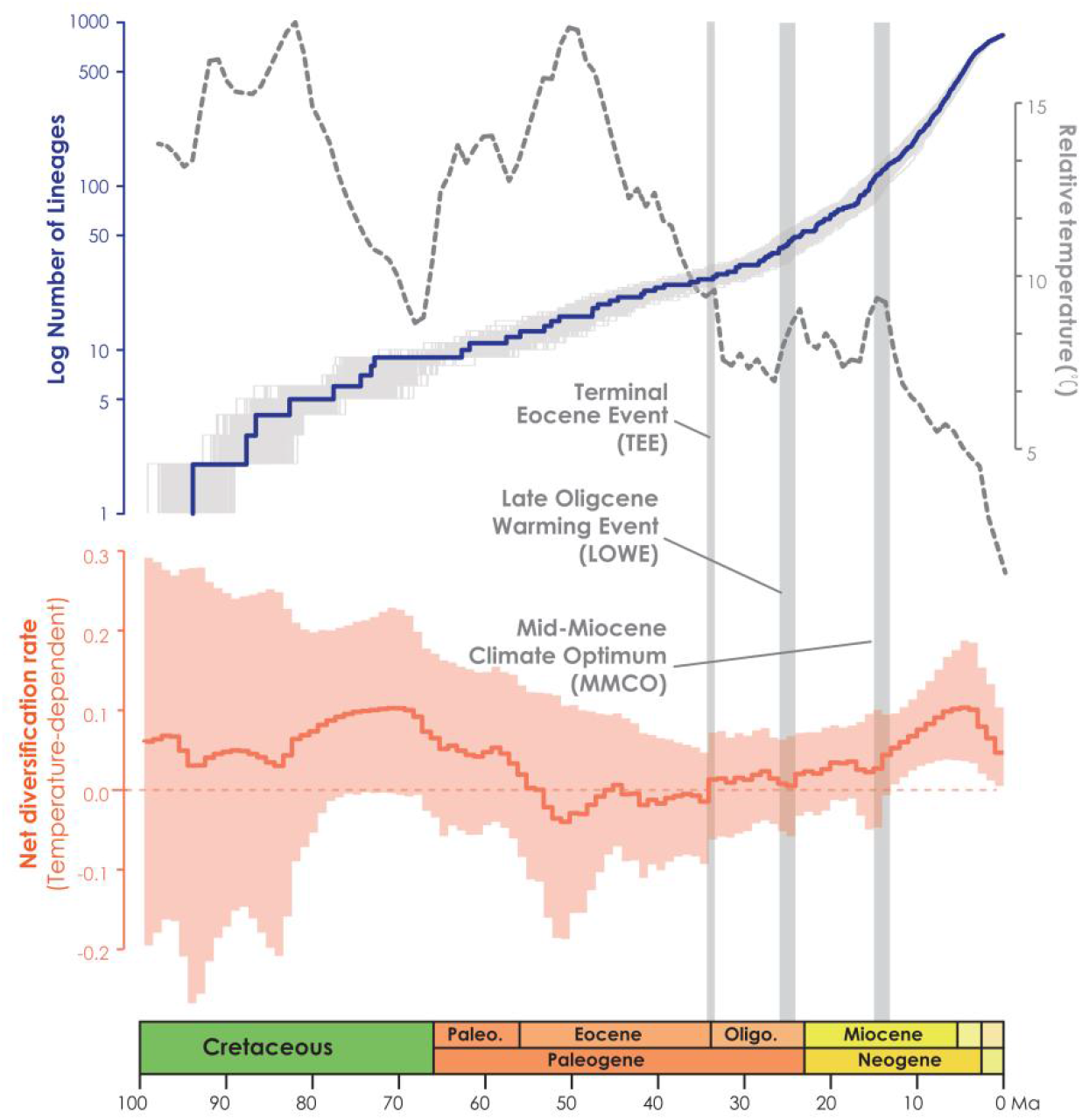
Summary figures include a global relative paleo-temperature plot (gray dotted line), lineage through time (LTT) plot (blue line), and net diversification rate through time plot in the function of temperature from RevBayes analysis for Annonaceae (orange plot: light orange area representing the credibility interval, and the continuous line representing the median rate).

### 3.2 Paleotemperature-dependent diversification

The temperature-dependent diversification rates plot based on the 835-taxon tree, represented by the orange line in Fig. 1, shows that the net diversification rate of Annonaceae has significantly exceeded zero from the late Oligocene. This suggests that speciation rate has accelerated under the influence of global temperature. In the RPANDA analysis (Table S1), the best-supported model with the lowest AICc value (5128.185, AICcω=1) indicates that both speciation and extinction rates are significantly positively correlated with paleotemperature, with positive alpha and positive beta values, respectively. The net diversification rates notably show a rapid increase from c. 25 Ma, as reported by Xue et al. (2020), which aligns with the trends observed in both time and temperature-dependent net diversification rates (Fig. 1).

### 3.3 Clade-dependent diversification

The best diversification rate shifts configuration estimation based on the 728-taxon tree reveals rate shifts in five clades (Fig. 2). The earliest rate shift occurred at the branch with the largest number of tribes within subfamily Annonoideae (excluding tribe Bocageeae) (Fig. 2: N1). This was followed by four subsequent rate shifts: in subfamily Malmeoideae (excluding tribe Piptostigmateae) (Fig. 2: N2); basal to the *Cyathocalyx*-*Drepananthus* clade within subfamily Ambavioideae (Fig. 2: N3); in *Guatteria* (subfamily Annonoideae tribe Guatterieae, excluding *Guatteria anomala*) (Fig. 2: N4); and basal to *Pseuduvaria* (subfamily Malmeoideae tribe Miliuseae) (Fig. 2: N5).

**Figure 2.**
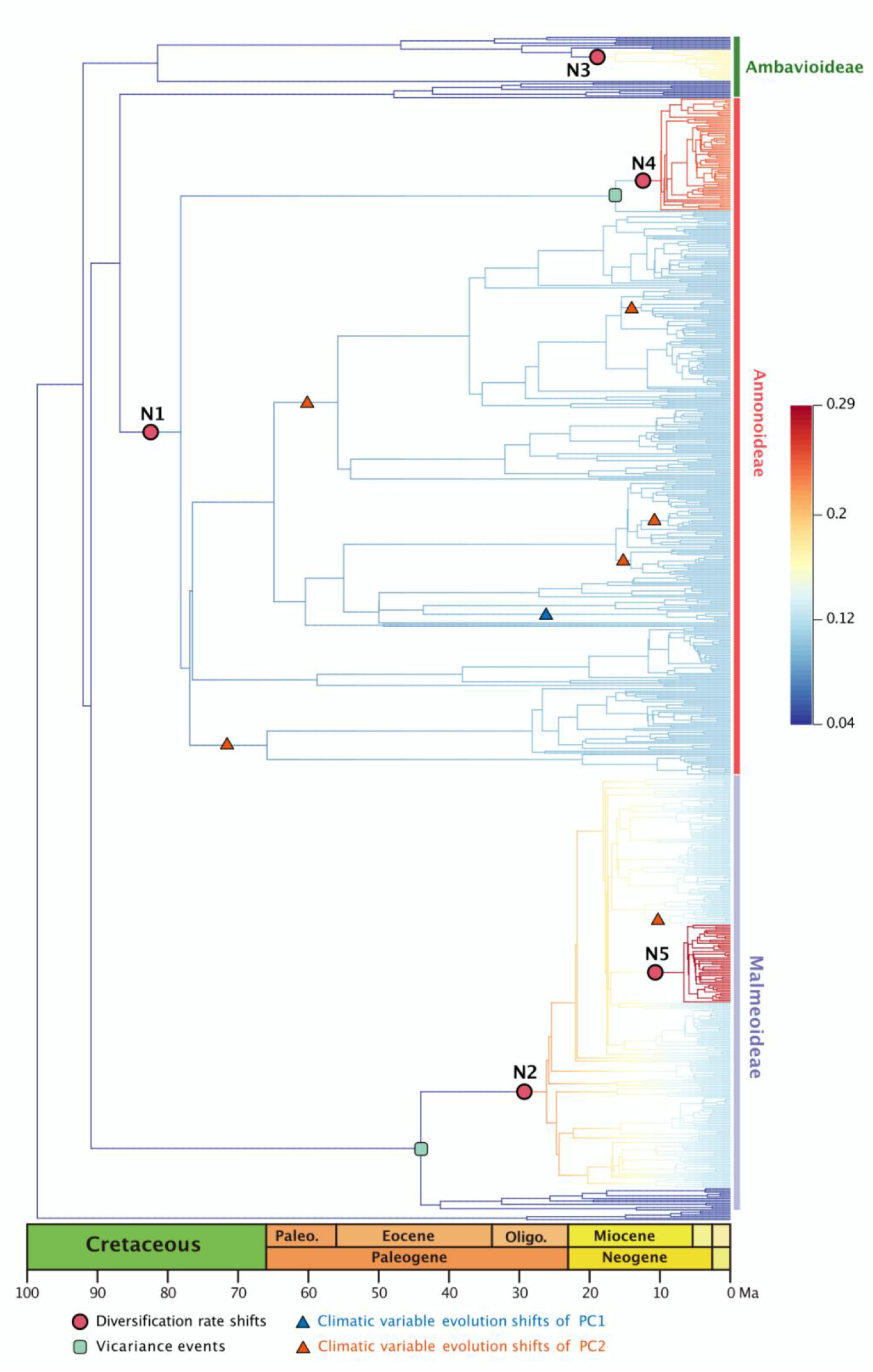
Best BAMM scenario, with diversification rate shift (shown in red dots) in Annonaceae per million, mapped positions of vicariance events (shown in the green square), climatic variable evolution shifts of PC1 (shown in blue triangle), climatic variable evolution shifts of PC2 (shown in red triangle). Branch colors indicating diversification rates (refer to the figure S3 for higher resolution)

### 3.4 Biogeographical events

Annonaceae are widely distributed in all six biogeographic regions (Figs 3 and S2). The best-fitting model for ancestral area reconstruction was DEC+j based on the 728-taxon tree. BSM analysis identified 1191 biogeographic events (Table S3d), with cladogenetic events 22% more frequent than anagenetic events. In situ speciation within an area accounted for 53.3% of events, vicariance for 4%, and dispersal for 42.5%, including cladogenetic (3.7%) and anagenetic (38.8%) dispersals (Table S3d). Areas B, C and F (eastern Southeast Asia, continental Africa, and South America) were common sources of dispersal events (Table S3). BSM analysis identified significant long-distance dispersal (LDD) events (Fig. 3), with prominent asymmetric floristic exchange observed (Table S3a): the magnitude between A and B (western and eastern Southeast Asia) is 2.4:1 (A to B: B to A), between E and F (North/Central and South America) is 2.7:1 (F to E: E to F).

**Figure 3.**
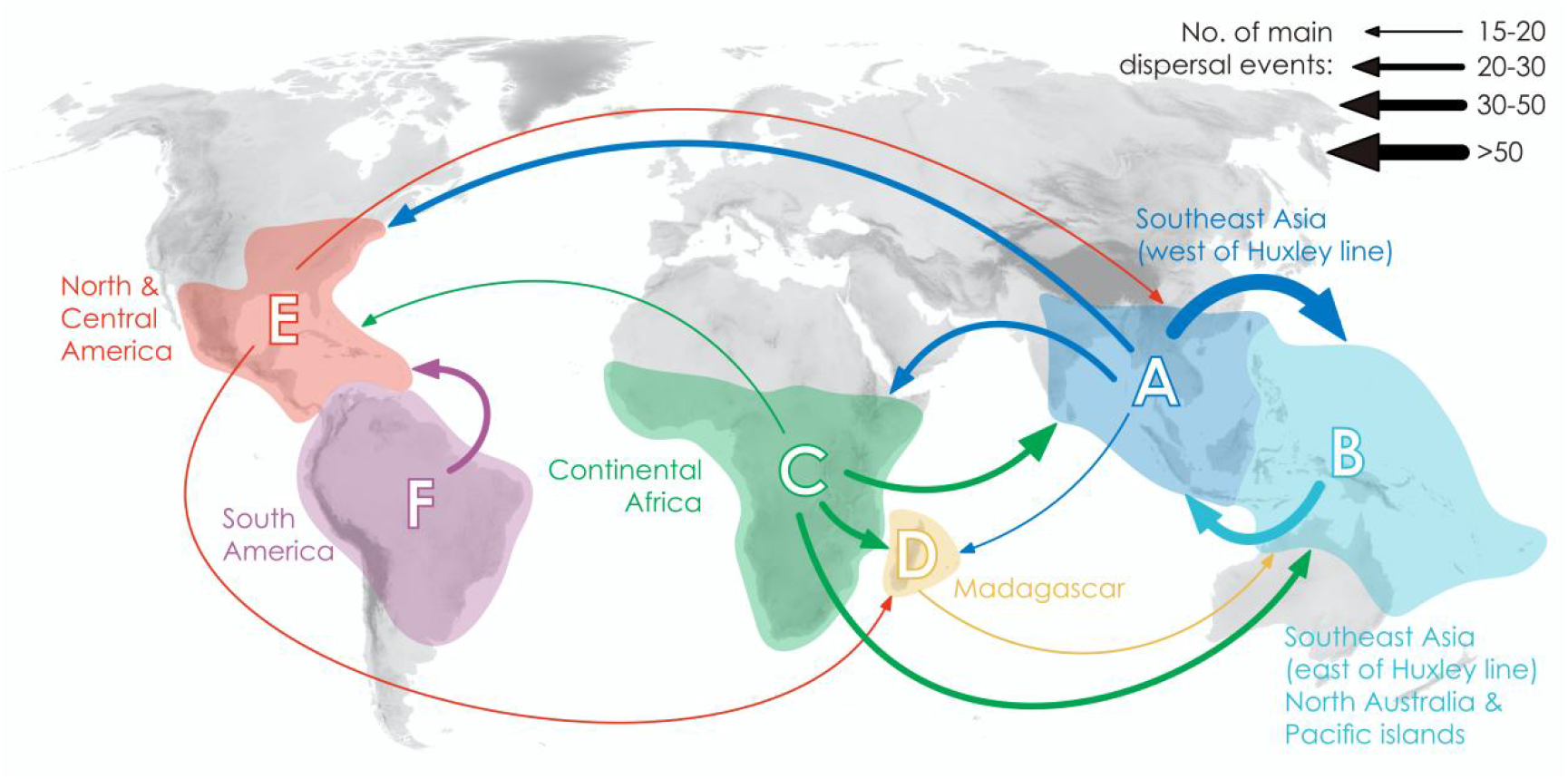
Delimitation of six geographical areas, and dispersal rate between different geographical areas (the thickness of the arrows represents the transition rates and transition directions).

Two of the four recent diversification rate shifts were recovered on the same node as reconstructed biogeographical events (Fig. 2): one was a vicariance event between *Guatteria* and its sister clade, while the other was a vicariance event in subfamily Malmeoideae: between tribe Bocageeae and the remaining tribes. The former vicariance event is inferred to have taken place between South America and North America, resulting in rapid diversification in the South American clade (Fig. 2). To further investigate the impact of geographical areas on Annonaceae diversification, GeoSSE model tests were conducted using South America as an individual area and combining all other areas. The best-fitting model was the GeoHiSSE model with one hidden trait and no range-dependent diversification, having the lowest AIC value (5115.024, AICω = 0.93) (Table S4). This suggests that the diversification rate for lineages is unrelated to endemicity in South America (Table S4). Consistent with this, other clades in South America do not exhibit fast diversification rate shifts. The HiSSE model test also indicated the presence of hidden factors that might impact Annonaceae diversification beyond the two defined areas (Table S6, AIC = 5252.590 with AICω = 1), consistent with a heterogeneous diversification process (above). Vicariance in the Malmeoideae was inferred to have occurred between Africa and other areas combined, suggesting that the clade underwent rapid diversification out of Africa after the event (Fig. 2). The GeoSSE model test showed that the best-fitting model was the GeoHiSSE model with one hidden trait and range-dependent diversification, having the lowest AIC value (5115.024, AICω = 1) (Table S5), indicating a higher diversification rate for lineages outside Africa (Table S5). The HiSSE model test again suggests the presence of hidden factors that might impact Annonaceae diversification beyond the two defined areas (Table S7, AIC = 4931.075 with AICω = 1).

### 3.5 Climatic niche evolution

In the PCA analysis based on the dataset of 728 species, PC1 primarily captures the annual stability and the minimum extreme temperature value during the driest and coldest period, whereas PC2 reflects precipitation during driest period (Table S8). Within PC1, a significant evolutionary niche shift (PP = 1) was identified at the *Asimina* crown node (Figs 2 and S5). *Asimina* primarily occupies temperate and subtropical habitats in eastern North America (Kral, 1960), with the reconstruction of the ancestral area of *Asimina* retrieved as East Asia (Fig. S2). It is worth noting that *Asimina* is the only Annonaceae genus fully adapted to freezing temperatures (Hormaza, 2014). In PC2, a total of six optimal evolutionary shifts (PP > 0.5) were observed, with five shifts occurring in subfamily Annonoideae and one in subfamily Malmeoideae (Figs 2 and S5). No biogeographical events or species diversification rate shifts were directly associated with these niche evolutionary shifts (Fig. 2). QuaSSE analyses furthermore revealed significant correlations between speciation rates and climatic variables (PC1 and PC2) (Tables S9 and S10). The timing plot (Fig. 4) notably displayed a rapid increase in species diversification rates around 25 Ma, followed by a sudden divergence in climatic niche values around 15 Ma, further suggesting a difference in timing between these two evolutionary rates.

**Figure 4.**
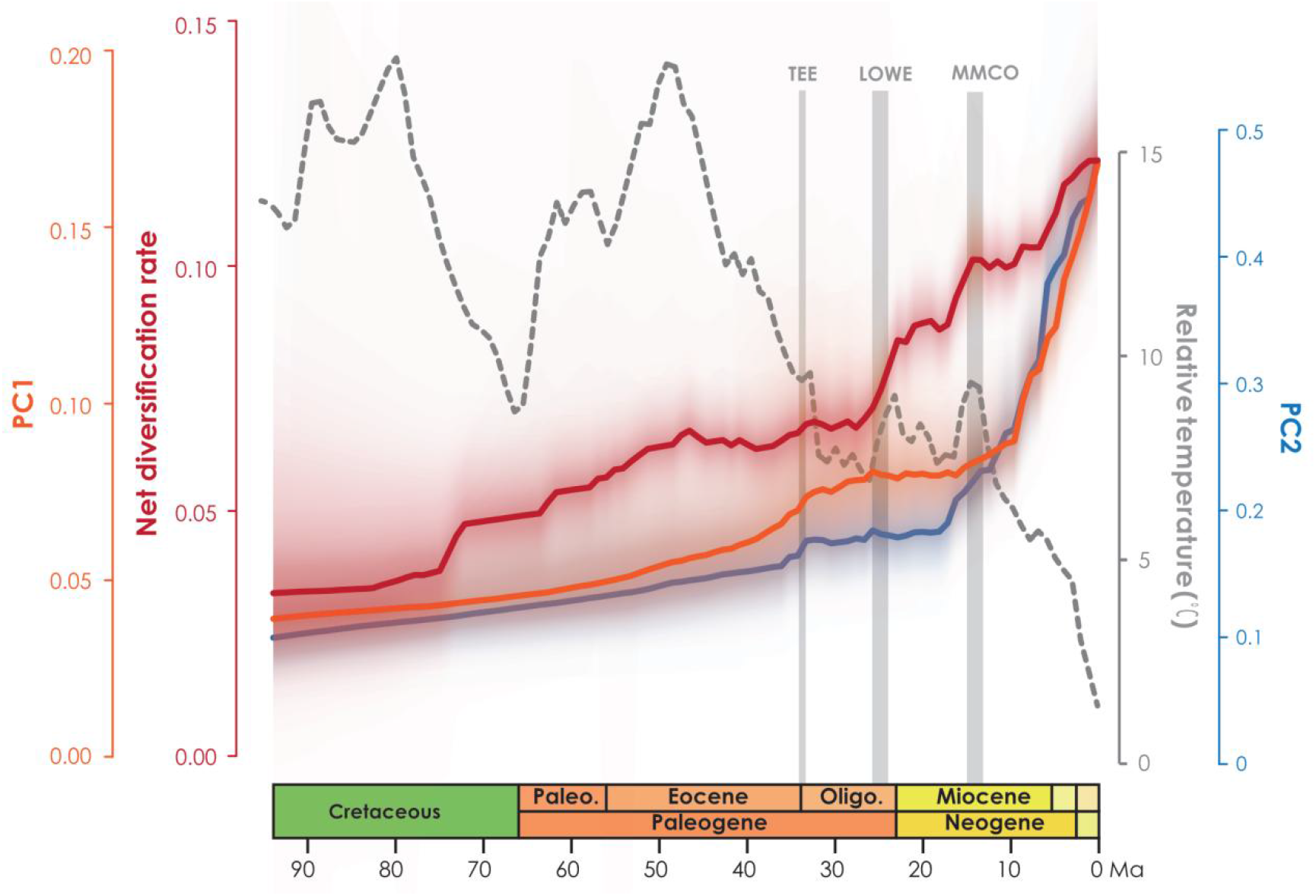
Net diversification rate (red line) and climatic PC1 and PC2 evolution rates over time (orange line and blue line, respectively).

Climatic niche reconstruction (Fig. S4) shows that most of the jumps with higher PP values in the reconstruction results of PC1 are adaptations to lower values (yellow to red), indicating that jumps in Annonaceae tend to be in response to increased climatic extremes (cooler and drier climate). In the ancestral reconstruction results of PC2, the jumps with larger PP values are likewise all towards lower values (yellow to orange), indicating that the clade of these jumps are in response to increasingly arid climatic extremes.

## 4. Discussion

### 4.1 Does global temperature drive the diversification in the Annonaceae?

Both family- and subfamily-wide LTT plots lend credence to our assumption: the recent diversification pattern of Annonaceae, characterized by fast acceleration since around 30 Ma, fits better with the recent cradle model. The museum model appears to be applicable during the earlier stages of diversification, and hence it is likely that the museum model transitioned into the recent cradle model (Figs 1 and S1). Our results challenge the previous assumption that a museum model of lineage accumulation is applicable to the Annonaceae (Couvreur et al., 2011), which could be attributed to a more comprehensive taxon sampling strategy (Sun et al. 2020b). Temperature-dependent analysis (Fig. 1) and RPANDA model tests (Table S1, AICω = 1) furthermore support the influence of global temperature on the accumulation of net diversification, which reinforce our hypothesis that global temperature change is a crucial driver for the recent diversification of Annonaceae. Diversification driven by global temperature is a common phenomenon in various plant lineages, such as *Oreocharis* (Gesneriaceae; Kong et al., 2022), *Primula* (Primulaceae; Smyčka et al., 2022), and rosids (Sun et al., 2020a).

The five identified diversification rate shifts in our study (Fig. 2) exhibit slight variations compared to those reported by Xue et al. (2020), possibly due to the exclusion of taxa without specific locality records in our analysis. However, not all of those diversification rate shifts can be directly attributed to global climate events, suggesting the involvement of other factors in macroevolutionary processes. In terms of specific clades within Annonaceae, the onset of high diversification rates of Malmeoideae is speculated to be related to the Terminal Eocene Event (TEE, ca. 34 Ma), which involved a sharp drop in global temperature and likely promoted vicariance events. Instead, the timing of diversification rate shifts in the *Gua****t****eria* and *Pseuduvaria* clades align with the end of Mid-Miocene Climatic Optimum (MMCO, ca. 16–14 Ma), the resultant sudden global cooling and drought forced species to encounter new ecological opportunities for speciation. Similar patterns have been observed in other plant lineages, such as *Hypericum* (Hypericaceae), in which massive mountain formation and climate cooling following the last thermal maximum in the Miocene stimulated global radiation (Nürk et al., 2013).

### 4.2 Do diversification rates reflect climate niche lability?

The plot of niche evolution over time (Fig. 4) reveals that the acceleration of climate niche shift (ca. 15 Ma) appears to be later than the rapid increase in the diversification rate of Annonaceae (ca. 25 Ma). The onset of species diversification around 25 Ma may have occurred in ancestral areas, where the “Late Oligocene Warming Event” (LOWE, ca. 26–24 Ma) provided optimal global climate conditions for tropical rainforest biome development, leading to expanded habitats that promoted species dispersal and exchange. The initial diversification may therefore reflect niche conservatism, as no rapid increase in niche evolutionary rate was observed before the mid-Miocene. Numerous studies have provided evidence that species dispersal enables them to evade competition, facilitating their adaptation to new environments (Burgess et al., 2016; Economou, 1991; Martorell & Martínez-López, 2014), suggesting the importance of geographical transitions in achieving niche vicariance (Rosenzweig, 1995; Gómez-Rodríguez et al., 2015). Similarly, the diversification process itself enables greater niche vicariance, leading to accelerated niche evolution. A lag between niche evolution and the rapid increase in diversification rate (Fig. 4) is uncommon but has also been documented in Saxifragales (Folk et al., 2019) and *Penstemon* (Plantaginaceae; Stone & Wolfe, 2021). Different timing patterns has been documented in other taxa. For instance, in *Hypericum* (Hypericaceae) niche evolution precedes the burst in diversification rate, with the early adaptation to drought tolerance enabling them to thrive during the sharp global temperature drop following the mid-Pliocene thermal optimum (Nürk et al., 2013). In contrast to the interpretation by Folk et al. (2019) and Stone & Wolfe (2021), however, our results indicate that diversification in the Annonaceae does not appear to be influenced by density-dependent factors, as the net diversification rate shows an increasing trend in general (Fig. 4), challenging the established concept of density-dependent diversification rate slowdown associated with the saturation of ecological niches (Morlon et al., 2011), but appears to be influenced by interspecific competition. The strong asymmetric floristic exchange observed in the Annonaceae (Fig. 3 and Table S3) supports our assumption that interspecific competition leads to species dispersing from areas with higher to lower species richness, and Areas A and F, which are old and species-rich regions, serve as common dispersal sources (Bacon et al., 2015; Crayn et al., 2015; Dupin et al., 2017). Our findings challenge the assumption that shifts in niche evolution directly dictate changes in diversification rate. Even though our QuaSSE model test results (Tables S9 and S10) suggest that the speciation rate is clearly correlated with PCs raw values, the niche evolutionary test show no direct correspondence with shifts in divergence rate (Fig. 2). Similar process might occur in other taxa: diversification in Terebinthaceae (Weeks et al., 2014) and phaseoloid legumes (Li et al., 2013) may not be driven by niche evolution as speculated; instead, it is possible that they exhibit a similar diversification pattern as Annonaceae. Our study therefore provides a novel perspective for understanding diversification across disparate taxa.

### 4.3 Are particular biogeographical events associated with shifts in diversification rate in the Annonaceae?

Compared with a large number of estimated biogeographical events, only two of five diversification rate shifts are directly associated with geographical events, indicating that geographical events do not always trigger diversification. Both mapped biogeographical events are vicariant, including the *Gua****t****eria*clade, which experienced rapid diversification in South America following a vicariance event. This finding is consistent with previous research by Erkens et al. (2007), which also supports the rapid diversification of *Guatteria* after its colonization of South America. The uplift of the Andes mountains in South America, particularly during the Late Miocene and early Pliocene, created opportunities for diversification by promoting physiographic heterogeneity, regional climate diversity, parallel geographic speciation, and adaptive radiation (Garzione et al., 2008; Mulch et al., 2010; Hoorn et al., 2010, Hoorn 2013; Luebert & Weigend, 2014; Lagomarsino et al., 2016; Mulch, 2016). Similar diversification patterns associated with Andean uplift have been observed in lineages of other plant families, such as the *Oxalis tuberosa* alliance (Oxalidaceae; Emshwiller & Doyle, 1998, 2002), *Lupinus*(Fabaceae; Hughes & Eastwood, 2006; Drummond et al., 2012), *Hypericum*(Hypericaceae; Nürk et al., 2013), and the Páramo clade in the Valerianaceae (Madriñán et al., 2013). Other intrinsic biotic factors of *Gua****t****eria*, such as pollen monads and absence of anther septation, have been reported to be related to high net diversification rates (Xue et al., 2020). In contrast to the *Gua****t****eria* clade, the reasons behind the diversification rate shift in the other clades are more complex. The second branch with diversification rate shift comprises 45 genera and spans five biogeographical regions, excluding Africa. Recent global climate change and frequent tectonic movements may have contributed to diversification rate shifts in regions outside of Africa (Maslin et al., 2005; Antonelli et al., 2018).

A similar pattern has also been reported in *Scleria* (Cyperaceae; Larridon et al., 2021), Adoxaceae and Valerianaceae (Moore & Donoghue, 2007) in which the relationship between dispersal and diversification is not a simple deterministic relationship, suggesting the involvement of key innovation factors. The results of the GeoSSE analysis did not provide conclusive evidence to support range-dependent diversification of Annonaceae (Tables S4 and S5), and HiSSE analysis also suggests the influence of geographic regions and other hidden traits on diversification (Tables S6 and S7). Geographical events undoubtedly play a crucial role in diversification, and asymmetric dispersal contributes to the niche evolution resulting from diversification, but our findings overall do not provide conclusive evidence for a direct association between range evolution and species diversification in Annonaceae. It is important to note that not all shifts in diversification rate can be solely attributed to biogeographical events, as they need to occur in the right location and timeframe for their influence to be significant. In the future, comprehensive taxon sampling as well as estimation of biogeographical events in extinct lineages are required to further test hypotheses regarding the correlation between dispersification and biological characteristics, and the interplay between range and niche evolution in diversification.

### 4.4 The future fate of pantropical plants following diversification

Our results also foreshadow the potential future fate of pantropical plant lineages following diversification, including ongoing expansion into temperate and cold regions. Shifts in niche evolution appear to be associated with increasing seasonal environments (Table S8 and Fig. S4), driven by species diversification. *Asimina*(Annonaceae) is a good example of a genus which experienced climatic niche shift following migration from tropical East Asia to temperate North America (Li et al., 2017), and following the evolution of protogynous flowering with an extended flowering period (Kral, 1960; Willson & Schemske, 1980; Lagrange et al., 1985; Norman & Clayton, 1986; Norman et al., 1992; Cox et al., 1998; Losada et al., 2017) and stigma secretions covering all stigmas (Losada et al., 2017), allowing *Asimina*to thrive in colder and drier habitats. A previous study of biotic correlations (Xue et al., 2020) also demonstrated that Annonaceae possess adaptive traits that allow them to thrive in diverse climatic conditions, indicating their potential for expansion beyond the tropics. Our study further highlights the capacity of Annonaceae to disperse over long distances, enabling them to reach and establish in new geographic areas. Future research exploring the interplay between niche lability and range evolution will further enhance our understanding of the dynamics of pantropical plant lineages.

## 5. Conclusions

Species diversification in the Annonaceae family initially followed the museum model, and shifts to a pattern corresponding to the recent cradle model from the late Oligocene. Global temperature has played a significant role in driving this diversification pattern. Our findings contradict the hypothesis that niche vicariance drives the diversification of Annonaceae: instead, species in the family tend to escape interspecific competition after diversification, resulting in later niche vicariance. While particular areas of niche space, particularly those reflecting greater abiotic challenge, are associated with diversification in Annonaceae, shifts in niche evolution do not accord with shifts in diversification rate. Our findings not only provide insights into the past diversification patterns of pantropical plant groups but also have implications for assessing their future fate of expansion into temperate regions. However, we have yet to address the interplay among key opportunities, key innovation, and species diversification. Future comprehensive analyses of range evolution are needed to further test this hypothesis.

## CRediT authorship contribution statement

**Weixi Li:** Conceptualization, Investigation, Formal analysis, Data curation, Writing – original draft. **Runxi Wang:** Formal analysis, Writing – review & editing. **Ming-Fai Liu:** Writing – review & editing. **Ryan A. Folk:** Supervision, Writing – review & editing. **Bine Xue:** Conceptualization, Supervision, Resources, Writing – review & editing. **Richard M.K. Saunders:** Conceptualization, Funding acquisition, Supervision, Resources, Project administration, Writing – review & editing.

## Acknowledgement

We thank Dr. Gaitan-Espitia, Juan Diego for comments on the manuscript, and Laura Wong for general technical support.

