## Supplemental Figure 1 for "Climatic and biogeographic processes underlying the diversification of the pantropical and early divergent angiosperm family Annonaceae"

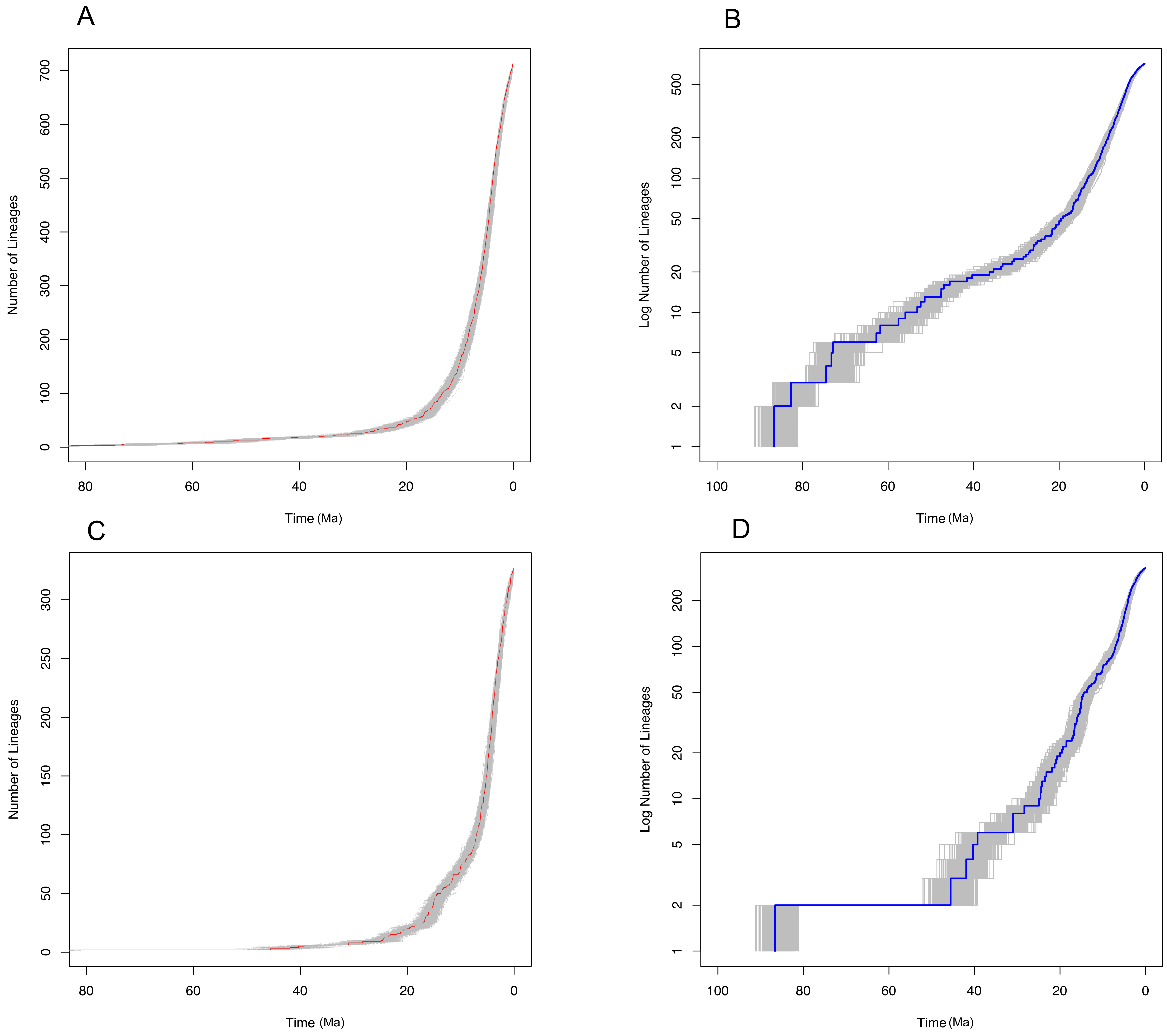


**Figure S1.** Number-of-lineage plots for subfamily Annonoideae (A) and subfamily Malmeoideae (C). Semilogarithmic mean lineage-through-time (LTT) plots for subfamily Annonoideae (B) and subfamily Malmeoideae (D). The solid red line and blue lines in the middle indicate the mean values derived from 1,000 posterior trees obtained from the Bayesian analysis. The gray area represents the upper and lower 95% confidence intervals.
