## Supplemental Figure 2 for "Climatic and biogeographic processes underlying the diversification of the pantropical and early divergent angiosperm family Annonaceae"

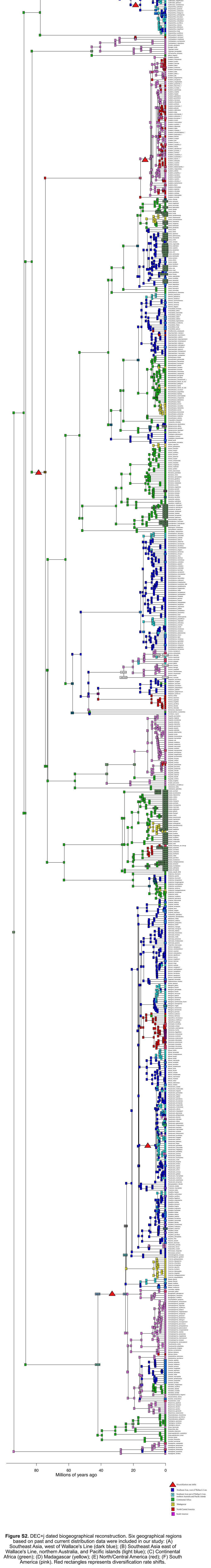


**Figure S2.** DEC+j dated biogeographical reconstruction. Six geographical regions based on past and current distribution data were included in our study: (A) Southeast Asia, west of Wallace's Line (dark blue); (B) Southeast Asia east of Wallace's Line, northern Australia, and Pacific islands (light blue); (C) Continental Africa (green); (D) Madagascar (yellow); (E) North/Central America (red); (F) South America (pink). Red rectangles represents diversification rate shifts.
