## Supplemental Figure 3 for "Climatic and biogeographic processes underlying the diversification of the pantropical and early divergent angiosperm family Annonaceae"

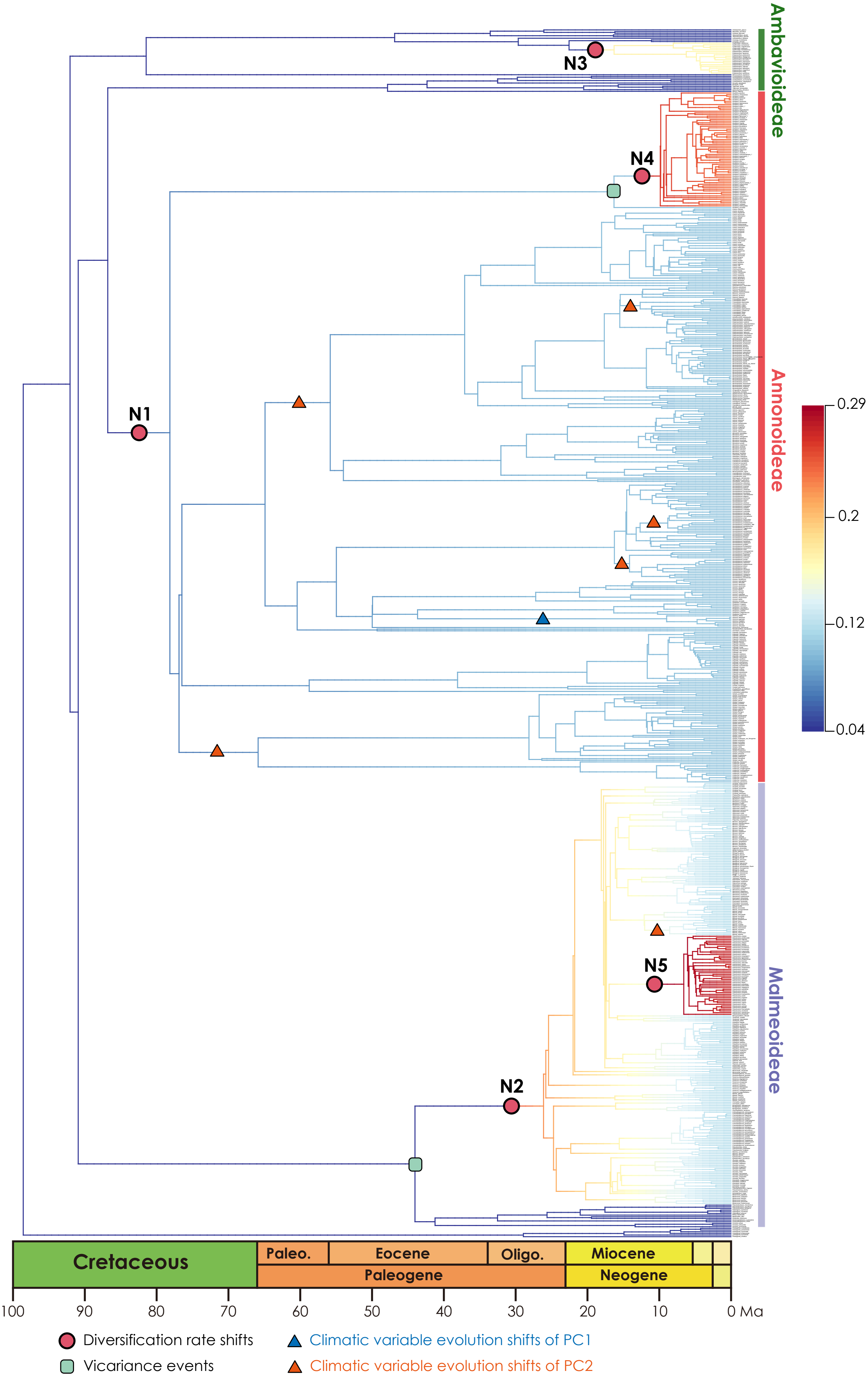


**Figure S3**. Best BAMM scenario, with diversification rate shift (shown in red dots) in Annonaceae per million, mapped positions of vicariance events (shown in the green square), climatic variable evolution shifts of PC1 (shown in blue triangle), climatic variable evolution shifts of PC2 (shown in red triangle). Branch colors indicating diversification rates.
