## Supplemental Tables for "Climatic and biogeographic processes underlying the diversification of the pantropical and early divergent angiosperm family Annonaceae"

**Table S1.** The constant, time-dependent, temperature-dependent and temperature-fluctuation dependent model test result by RPANDA. The best model is in bold.

| No | Models | Rate_variation | NP | logL | AICc | AICc_ω | Lambda | Alpha | Mu | Beta |
|---|---|---|---|---|---|---|---|---|---|---|
| 1 | BCST | constant | 1 | -2699.545 | 5401.095 | 5.47453E-60 | 0.1848 | NA | NA | NA |
| 2 | BCSTDCST | constant | 2 | -2644.069 | 5292.152 | 2.48324E-36 | 0.3499 | NA | 0.2452 | NA |
| 3 | BTimeVar_EXPO | exponential | 2 | -2589.331 | 5182.676 | 1.47038E-12 | 0.267 | -0.0405 | NA | NA |
| 4 | BTimeVarDCST_EXPO | exponential | 3 | -2589.351 | 5184.73 | 5.26513E-13 | 0.2653 | -0.0398 | 0 | NA |
| 5 | BCSTDTimeVar_EXPO | exponential | 3 | -2616.97 | 5239.968 | 5.32881E-25 | 0.3072 | NA | 0.1554 | 0.0113 |
| 6 | BTimeVarDTimeVar_EXPO | exponential | 4 | -2589.331 | 5186.71 | 1.9564E-13 | 0.267 | -0.0404 | 0 | -0.0289 |
| 7 | BTimeVar_LIN | linear | 2 | -2597.896 | 5199.807 | 2.80208E-16 | 0.2368 | -0.0048 | NA | NA |
| 8 | BTimeVarDCST_LIN | linear | 3 | -2593.68 | 5193.389 | 6.93635E-15 | 0.2633 | -0.0053 | 0.0343 | NA |
| 9 | BCSTDTimeVar_LIN | linear | 3 | -2609.253 | 5224.534 | 1.19696E-21 | 0.2714 | NA | 0.0883 | 0.0034 |
| 10 | BTimeVarDTimeVar_LIN | linear | 4 | -2591.203 | 5190.454 | 3.00925E-14 | 0.2439 | -0.005 | 9.00E-04 | 9.00E-04 |
| 11 | BTempVar_EXPO | exponential | 2 | -2630.478 | 5264.97 | 1.98388E-30 | 0.3519 | -0.1228 | NA | NA |
| 12 | BTempVarDCST_EXPO | exponential | 3 | -2621.958 | 5249.944 | 3.63387E-27 | 0.3617 | -0.0666 | 0.1047 | NA |
| 13 | BCSTDTempVar_EXPO | exponential | 3 | -2608.211 | 5222.451 | 3.39155E-21 | 0.2877 | NA | 0.0902 | 0.0856 |
| 14 | BTempVarDTempVar_EXPO | exponential | 4 | -2592.527 | 5193.101 | 8.01069E-15 | 0.2184 | 0.2354 | 0.209 | 0.2384 |
| 15 | BTempVar_LIN | linear | 2 | -2622.603 | 5249.221 | 5.21635E-27 | 0.3139 | -0.0228 | NA | NA |
| 16 | BTempVarDCST_LIN | linear | 3 | -2618.659 | 5243.347 | 9.83761E-26 | 0.3339 | -0.0156 | 0.0931 | NA |
| 17 | BCSTDTempVar_LIN | linear | 3 | -2597.134 | 5200.298 | 2.1921E-16 | 0.244 | NA | 0.0389 | 0.0224 |
| **18** | **BTempVarDTempVar_LIN** | **linear** | **4** | **-2560.069** | **5128.185** | 1 | **0.0556** | **0.133** | **0.1506** | **0.1395** |

Note: B: birth rate; D: death rate; Var: vary; CST: constant; EXPO:Exponentially; LIN: linearly; Tem: temperature.

**Table S2.** BioGeoBEARS models comparison based on 728 species. Likelihood, number of parameters, parameters (d, e and j) and AICc are shown. The best model is in bold.

| | LnL | numparams | d | e | j | AIC | AIC_ω | AICc | AICc_ω |
|---|---|---|---|---|---|---|---|---|---|
| DEC | -1677.741209 | 2 | 0.013211371 | 2.00E-08 | 0 | 3359.482419 | 1.28E-187 | 3359.49897 | 1.29E-187 |
| **DEC+J** | **-1651.534686** | **3** | **0.012447941** | **1.00E-12** | **0.007894815** | **3309.069372** | **1.13E-176** | **3309.102521** | **1.13E-176** |
| DIVALIKE | -1751.859357 | 2 | 0.014883572 | 0.000227872 | 0 | 3507.718714 | 8.26E-220 | 3507.735266 | 8.32E-220 |
| DIVALIKE+J | -1715.379294 | 3 | 0.013672579 | 1.00E-12 | 0.007976148 | 3436.758588 | 2.12E-204 | 3436.791737 | 2.12E-204 |

**Table S3.** Counts of biogeographical events (and standard deviations, SD) between the different considered regions inferred by the BSM analysis under the DEC+j model. Notes: (A) Southeast Asia, west of Wallace's Line; (B) Southeast Asia east of Wallace's Line, northern Australia and Pacific islands ; (C) Continental Africa; (D) Madagascar; (E) North/Central America; (F) South America.

**(a)** Summary of the number of dispersal events (and standard deviations, SD). The main movements are in bold.

| | | To | | | | | |
|---|---|---|---|---|---|---|---|
| | | A | B | C | D | E | F |
| From | A | 0<br>(0) | **82.72<br>(8)** | **22.88<br>(4.14)** | 19.76<br>(4.64) | **26.84<br>(4.55)** | 8.98<br>(1.68) |
| | B | **34.16<br>(7.63)** | 0<br>(0) | 13.54<br>(3.7) | 14.18<br>(3.29) | 13.92<br>(4.51) | 3.2<br>(1.21) |
| | C | **27.12<br>(4.73)** | **20.84<br>(4.29)** | 0<br>(0) | **24.8<br>(4.97)** | 19.8<br>(5.35) | 4.74<br>(1.48) |
| | D | 12.94<br>(3.6) | 15.14<br>(4.18) | 12.94<br>(3.55) | 0<br>(0) | 14.16<br>(4.09) | 0.5<br>(0.68) |
| | E | 18.54<br>(4.5) | 14.26<br>(3.97) | 13.62<br>(4.41) | 15.06<br>(4.71) | 0<br>(0) | 10.46<br>(2.52) |
| | F | 3.6<br>(1.53) | 2.8<br>(1.47) | 2.9<br>(1.59) | 1.34<br>(0.85) | **28.2<br>(2.69)** | 0<br>(0) |

**(b)** Summary of the number of anagenetic dispersal (range expansion) events (and standard deviations, SD).

| | | To | | | | | |
|---|---|---|---|---|---|---|---|
| | | A | B | C | D | E | F |
| From | A | 0<br>(0) | 73.82<br>(7.58) | 20.74<br>(4.11) | 19.32<br>(4.6) | 24.18<br>(5.01) | 7.46<br>(1.42) |
| | B | 28.4<br>(7.23) | 0<br>(0) | 13.36<br>(3.7) | 14<br>(3.23) | 13.8<br>(4.53) | 1.72<br>(1.11) |

| | | | | | | | |
|---|---|---|---|---|---|---|---|
| | C | 25.92 (4.56) | 20.52 (4.27) | 0 (0) | 20.9 (4.95) | 19.56 (5.38) | 3.56 (1.45) |
| | D | 12.7 (3.59) | 15.06 (4.2) | 12.8 (3.51) | 0 (0) | 14.1 (4.03) | 0.5 (0.68) |
| | E | 18.28 (4.42) | 14.18 (3.97) | 13.12 (4.37) | 15.02 (4.64) | 0 (0) | 8.44 (2.08) |
| | F | 2.66 (1.35) | 2.1 (1.39) | 2.24 (1.38) | 1.34 (0.85) | 20.28 (2.31) | 0 (0) |

**(c)** Summary of the number of cladogenetic dispersal (founder) events (and standard deviations, SD).

| | | To | | | | | |
|---|---|---|---|---|---|---|---|
| | | A | B | C | D | E | F |
| **From** | A | 0 (0) | 8.9 (2.28) | 2.14 (0.99) | 0.44 (0.64) | 2.66 (1.04) | 1.52 (1.11) |
| | B | 5.76 (1.53) | 0 (0) | 0.18 (0.44) | 0.18 (0.39) | 0.12 (0.33) | 1.48 (0.89) |
| | C | 1.2 (1.05) | 0.32 (0.59) | 0 (0) | 3.9 (1.11) | 0.24 (0.43) | 1.18 (0.83) |
| | D | 0.24 (0.48) | 0.08 (0.27) | 0.14 (0.35) | 0 (0) | 0.06 (0.24) | 0 (0) |
| | E | 0.26 (0.49) | 0.08 (0.27) | 0.5 (0.61) | 0.04 (0.2) | 0 (0) | 2.02 (1.27) |
| | F | 0.94 (0.96) | 0.7 (0.76) | 0.66 (0.77) | 0 (0) | 7.92 (2.14) | 0 (0) |

(d) Summary of the number of biogeography events in Annonaceae.

| **Process** | **Type of event** | **Cladogenetic/ Anagenetic event** | **Mean (SD)** | **Percentage** |
|---|---|---|---|---|
| Dispersal | Founder events | Cladogenetic | 43.86(4.85) | 3.7 |
| | Range expansions | Anagenetic | 460(4.91) | 38.8 |
| Vicariance | Vicariance | Cladogenetic | 46.98(4.97) | 4.0 |
| Speciation within area | Narrow sympatry | Cladogenetic | 532.7(9.77) | 44.8 |
| | Speciation subset | Cladogenetic | 103.5(11.15) | 8.7 |
| Total | | Cladogenetic | 727(0) | 61.2 |
| | | Anagenetic | 460(4.91) | 38.8 |
| | | | 1187(4.91) | 100 |

**Table S4.** Model test result of GeoSSE: the individual area south America was treated as state 1, and all other areas combined was treated as state 2.

| | loglik | AIC | AIC ω |
|---|---|---|---|
| 1 | -2643.067 | 5292.134 | 3.237131e-39 |
| 2 | -2642.871 | 5295.742 | 5.329479e-40 |
| **3** | **-2552.512** | **5115.024** | **9.316230e-01** |
| 4 | -2549.124 | 5120.247 | 6.837700e-02 |

**Table S5.** Model test result of GeoSSE: individual area Africa was treated as state 1, and all other areas combined was treated as state 2.  The best model is in bold.

| | loglik | AIC | AIC ω |
|---|---|---|---|
| 1 | -2769.877 | 5545.753 | 2.189466e-64 |
| 2 | -2727.003 | 5464.007 | 1.234131e-46 |
| 3 | -2676.797 | 5363.595 | 7.862041e-25 |
| **4** | **-2615.295** | **5252.590** | 1 |

**Table S6.** Model test result of HiSSE: south America was treated as state 0, and elsewhere was treated as state 1

| | **loglik** | **AIC** | **AIC ω** |
|---|---|---|---|
| bisse null | -2769.877 | 5545.753 | 2.1901E-64 |
| hisse null | -2727.003 | 5464.007 | 1.23419E-46 |
| habitat dependent | -2676.797 | 5363.595 | 7.86268E-25 |
| **habitat with hidden state** | **-2615.295** | **5252.590** | 1 |

**Table S7.** Model test result of HiSSE: Africa was treated as state 0, and elsewhere was treated as state 1. The best model is in bold.

| | loglik | AIC | AIC ω |
|---|---|---|---|

| | | | |
|---|---|---|---|
| bisse null | -2558.748 | 5125.495 | 6.05666E-43 |
| hisse null | -2507.988 | 5025.975 | 2.02406E-60 |
| habitat dependent | -2552.383 | 5116.765 | 4.76353E-41 |
| **habitat with hidden state** | **-2455.537** | **4931.075** | 1 |

**Table S8.** The quality of representation of 19 climatic variables across all dimensions. The variables that are well represented (with absolute value > 0.8) by PC1 and PC2 are in bold.

| | Bioclimatic variables | PC 1 | PC 2 | PC 3 | PC 4 | PC 5 |
|---|---|---|---|---|---|---|
| bio1_log | Annual Mean Temperature | **0.959** | -0.186 | 0.155 | -0.027 | -0.003 |
| bio2_log | Mean Diurnal Range (Mean of monthly (max temp - min temp)) | -0.621 | -0.235 | 0.152 | 0.077 | 0.571 |
| bio3_log | Isothermality (BIO2/BIO7) (×100) | **0.899** | 0.149 | 0.000 | -0.298 | 0.109 |
| bio4_log | Temperature Seasonality (standard deviation ×100) | **-0.866** | -0.260 | 0.023 | 0.292 | -0.058 |
| bio5_log | Max Temperature of Warmest Month | 0.220 | -0.605 | 0.681 | 0.299 | 0.109 |
| bio6_log | Min Temperature of Coldest Month | **0.975** | -0.064 | 0.043 | -0.131 | -0.018 |
| bio7_log | Temperature Annual Range (BIO5-BIO6) | **-0.920** | -0.210 | 0.066 | 0.249 | 0.170 |
| bio8_log | Mean Temperature of Wettest Quarter | 0.629 | -0.240 | 0.126 | 0.171 | -0.470 |
| bio9_log | Mean Temperature of Driest Quarter | **0.880** | -0.089 | 0.150 | -0.104 | 0.180 |
| bio10_log | Mean Temperature of Warmest Quarter | 0.564 | -0.445 | 0.577 | 0.298 | -0.144 |
| bio11_log | Mean Temperature of Coldest Quarter | **0.968** | -0.106 | 0.051 | -0.112 | 0.041 |
| bio12_log | Annual Precipitation | 0.686 | 0.590 | -0.062 | 0.284 | 0.192 |
| bio13_log | Precipitation of Wettest Month | **0.883** | 0.118 | -0.206 | 0.245 | 0.189 |
| bio14_log | Precipitation of Driest Month | -0.181 | **0.904** | 0.291 | 0.082 | -0.048 |
| bio15_log | Precipitation Seasonality (Coefficient of Variation) | 0.614 | -0.618 | -0.294 | -0.014 | 0.082 |
| bio16_log | Precipitation of Wettest Quarter | **0.863** | 0.170 | -0.197 | 0.281 | 0.225 |
| bio17_log | Precipitation of Driest Quarter | -0.126 | **0.922** | 0.299 | 0.063 | -0.074 |
| bio18_log | Precipitation of Warmest Quarter | 0.370 | 0.446 | -0.411 | 0.571 | -0.104 |
| bio19_log | Precipitation of Coldest Quarter | 0.134 | 0.726 | 0.499 | -0.178 | 0.113 |

**Table S9.** QuaSSE analyses result: Anova comparison between different models: Degrees of freedom, loglikelihood, AIC, ChiSq and P value are indicated for PC1 value and each of the five models. The best model is in bold.

| | Df | lnLik | AIC | AIC ω | ChiSq | Pr(>\|Chi\|) |
|---|---|---|---|---|---|---|
| minimal | 3 | -3357.055 | 6720.109 | 1.48612E-49 | NA | NA |
| linear | 4 | -3316.857 | 6641.714 | 1.56788E-32 | 80.395 | 0 |
| sigmoidal | 6 | -3294.951 | 6601.903 | 6.92434E-24 | 124.206 | 0 |
| drift.linear | 5 | -3254.053 | 6518.105 | 1.08809E-05 | 206.003 | 0 |
| **drift.sigmoidal** | 7 | -3240.624 | 6495.248 | 1 | **232.861** | **0** |

**Table S10.** QuaSSE analyses result: Anova comparison between different models: Degrees of freedom, loglikelihood, AIC, ChiSq and P value are indicated for PC2 value and each of the five models.The best model is in bold.

| | **Df** | **lnLik** | **AIC** | **AIC ω** | **ChiSq** | **Pr(>\|Chi\|)** |
|---|---|---|---|---|---|---|
| minimal | 3 | -3755.212 | 7516.424 | 8.52071E-25 | NA | NA |
| **linear** | **4** | **-3699.388** | **7406.776** | **0.550** | **111.648** | **0** |
| sigmoidal | 6 | -3701.308 | 7414.616 | 0.011 | 107.809 | 0 |
| **drift.linear** | **5** | **-3698.636** | **7407.273** | **0.429** | **113.151** | **0** |
| drift.sigmoidal | 7 | -3700.378 | 7414.756 | 0.010 | 109.668 | 0 |
